# Inability to make facial expressions dampens emotion perception

**DOI:** 10.1101/2022.10.11.510399

**Authors:** Shruti Japee, Jessica Jordan, Judith Licht, Savannah Lokey, Moebius Syndrome Research Consortium, Gang Chen, Joseph Snow, Ethylin Wang Jabs, Bryn D Webb, Elizabeth C Engle, Irini Manoli, Chris Baker, Leslie G. Ungerleider

## Abstract

Humans rely heavily on facial expressions for social communication to convey their thoughts and emotions and to understand them in others. One prominent but controversial view is that humans learn to recognize the significance of facial expressions by mimicking the expressions of others. This view predicts that an inability to make facial expressions (e.g., facial paralysis) would result in reduced perceptual sensitivity to others’ facial expressions. To test this hypothesis, we developed a diverse battery of sensitive emotion recognition tasks to characterize emotion perception in individuals with Moebius Syndrome (MBS), a congenital neurological disorder that causes facial palsy. Using computer-based emotion detection tasks we systematically assessed emotion perception thresholds for static and dynamic face and body expressions. We found that while MBS individuals were able to perform challenging perceptual control tasks, they were less efficient at extracting emotion from facial expressions, compared to matched controls. Exploratory analyses of fMRI data from a small group of MBS participants suggested potentially reduced engagement of the amygdala in MBS participants during expression processing relative to matched controls. Collectively, these results support the role of facial mimicry and consequent facial feedback and motor experience in the perception of others’ expressions.

## INTRODUCTION

Long before children are able to communicate verbally, their facial expressions convey their thoughts and emotions. Babies learn to portray their needs via the language of facial expressions from the first appearance of the social smile at about two months^1^. In the same way that babies learn the names of people and objects around them, they also learn the motor sequences for smiling and making other facial expressions during development^2,3^. Facial mimicry, facial feedback and motor experience, are considered to be three critical components of how humans learn to make facial expressions, feel the associated emotions, and perceive the same in others. For example, humans are thought to learn to express emotions via facial mimicry^4^ which can be thought of as the human version of *monkey see, monkey do*^*5-8*^. Thus, as an infant sees its parent smile, the visual cues of the observed motion are converted into motor actions of the baby’s own facial muscles^9,10^. Further, according to the facial feedback hypothesis, this motor action in turn provides proprioceptive feedback to the brain, which helps encode the sensation of the motor action^11,12^. Eventually, babies learn the emotional tags of these motor actions as ‘happiness, ‘sadness’, ‘anger’, etc. and associate them with the corresponding mental and physiological states. In addition to facial mimicry and facial feedback, motor experience is also thought to influence perception, such that expertise in performing a particular action, improves one’s perception of the same action in others. For example, professional actors have been shown to be better at explicit recognition of facial expressions^13^, while dancers who have expertise portraying emotion with their bodies, tend to be better at perceiving portrayed body expressions^14,15^. Consequently, facial movement could be considered a necessary condition for efficient facial motion perception and its absence could lead to impairment in the perception of facial expressions.

In this study, we investigated the role that facial movement, and thus facial mimicry, facial feedback, and motor experience have on emotion perception. We achieved this with a comprehensive battery of emotion detection tasks in a group of individuals with Moebius Syndrome (MBS). MBS is a rare congenital disorder characterized by limited lateral gaze and non-progressive facial paralysis due to underdevelopment or absence of the VI^th^ and VII^th^ cranial nerves. As a result, individuals with MBS are unable to smile, frown, grimace, or make other facial expressions from birth. Thus, this condition provides an ideal model to investigate how the inability to express emotions with one’s face, affects the ability to process and recognize emotions expressed by others.

The few prior studies of individuals with MBS have produced conflicting results, with some studies concluding that the inability to make facial expressions does not impact emotion perception^16-18^, while others have reported deficits in emotion recognition in individuals with MBS^19-22^. However, these prior studies had critical limitations. First, some studies^16,17^ did not include appropriate control tasks to account for differences in vision and general perceptual difficulties in MBS due to limited abduction of the eyes (i.e., inability to move eyes laterally because of the VI^th^ nerve palsy). Second, some of the previous studies used only full-blown facial expressions in the emotion recognition tasks^17^, which limit their sensitivity. Since individuals with MBS do not routinely report major difficulties with social communication, we expect that any impairment in emotion recognition would be subtle and require sensitive measurement methods. Third, none of the previous studies has examined the neural correlates of emotion processing in individuals with MBS and how they differ from healthy controls.

To address these issues, our study had three goals. First, we sought to systematically characterize emotion processing in MBS in the context of carefully designed perceptual control tasks. We hypothesized that if MBS participants could perform difficult non-emotional perceptual tasks similar to controls, but not emotional perceptual tasks, then this would provide strong evidence that emotion processing is affected in MBS. Second, we used a battery of sensitive emotion detection tasks to measure individual psychometric functions and detection thresholds. We did this by having participants detect emotion in images and videos containing varying levels of facial expressions and determining their perceptual sensitivity to emotional expressions by computing their thresholds for 50% detection accuracy. We hypothesized that while MBS participants may show no difference in recognition accuracy for full-blown facial expressions, we would see subtle differences in emotion detection thresholds in MBS compared to controls, since the inability to make facial expressions would likely affect their sensitivity to low levels of emotional information. Third, we used functional magnetic resonance imaging (fMRI) to explore whether individuals with MBS engage the same underlying neural circuitry for emotion processing, and to the same extent, as healthy controls. To do this, we used task-based fMRI in a small group of participants to delineate the neural circuits involved in emotional expression processing in MBS, and how they differ from brain networks normally engaged in the task.

## RESULTS

To systematically characterize emotion processing in MBS, we had individuals with MBS and a group of age- and gender-matched controls complete a comprehensive series of computer-based emotion detection and recognition tasks. A small subset of participants also completed task-based fMRI scans.

### Impaired detection of emotion from static facial expressions

First, we characterized emotion detection thresholds for happy and fearful faces in a *static facial expression task* by presenting morphs that contained different levels of the two facial expressions (Figures 1A and 1B). Briefly, participants were shown happy and fearful morphs in separate runs and asked to indicate if they thought the face was happy or neutral, or fearful or neutral, respectively, during *emotion detection task* runs, and to indicate if the mouth was open or closed during *feature detection control task* runs. The latter served as a critical perceptual control task to measure participants’ ability to process individual facial features. Morph levels yielding 50% detection accuracy were estimated as a percentage of expression information (0% morph level representing neutral expression and 100% morph level representing full-blown happy or fearful expression) by building the psychometric curves for each participant and each emotion, for each task. Overall, MBS individuals exhibited higher emotion detection thresholds than control participants, while feature detection thresholds did not differ between the two groups (Figure 1C). A mixed-effects beta regression analysis of threshold values showed the expected main effects of Task (Z = 6.80, p < 1.02 × 10^−11^) and Expression (Z = 8.70, p < 2.0 × 10^−16^), but more importantly a main effect of Group (Z = 3.92, p < 8.72 × 10^−5^), a two-way Task x Group (Z = 3.54, p < 0.0004) interaction, and a three-way Task x Expression x Group interaction (Z = 2.23, p < 0.025). Post-hoc Bonferroni-corrected between-group comparisons showed that for both happy and fearful face morphs, emotion detection thresholds for the *emotion task* were significantly higher (happy: t_124_ = 3.07, p < 0.003; fearful: t_124_ = 7.65, p < 2.02 × 10^−14^) for MBS individuals compared to controls (happy -- MBS: 30.3 ± 6.7% SD, Controls: 24.2 ± 5.6% SD; fearful -- MBS: 45.7 ± 13.9% SD, Controls: 29.2 ± 6.5% SD). By contrast, feature detection (open or closed mouth) thresholds for the *control task* were similar between the two groups for both happy (t_124_ = 1.64, p > 0.10; 50% thresholds for MBS: 23.2 ± 5.6% SD, Controls: 20.1 ± 5.2% SD) and fearful expressions (t_124_ = 1.39, p > 0.17; MBS: 30.3 ± 6.7% SD, Controls: 27.8 ± 6.4% SD).

**Figure 1.**
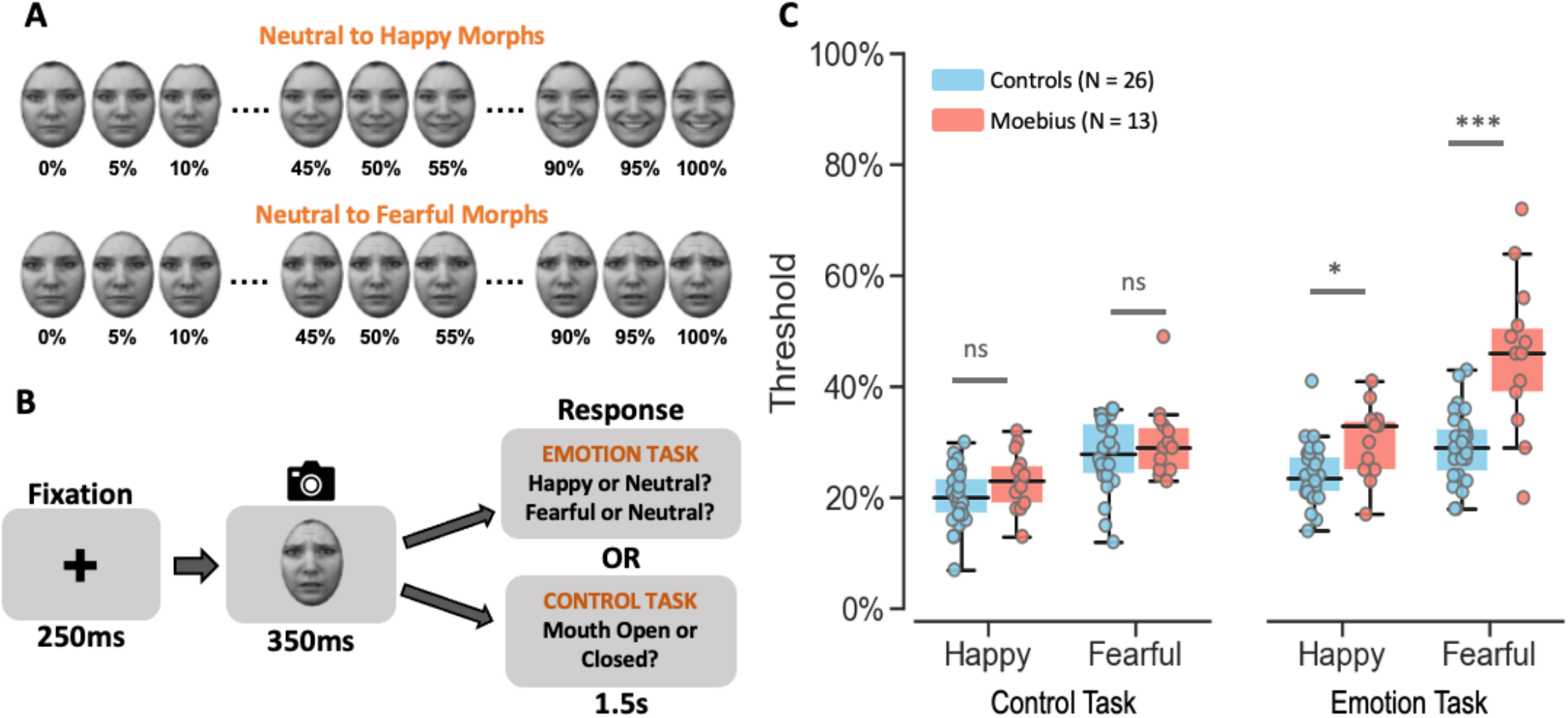
Static Facial Expression Task. **A**. Illustration of how stimuli were created by morphing a neutral face to its corresponding happy or fearful face in increments of 5% yielding 21 images for each emotion from 100% neutral to 100% happy or fearful. **B**. Depiction of the trial structure – each trial began with a 250ms fixation cross, followed by the happy or fearful morph image (in separate runs) for 350ms, and a 1.5s response window during which participants were to indicate if they thought the face was happy or neutral (or fearful or neutral in separate runs) during *emotion task* runs, or whether the mouth was open or closed during *control task* runs. **C**. Box and whisker plots showing the thresholds for each task for control participants in blue and MBS individuals in red. MBS individuals had similar thresholds as control participants on the *control task* but higher thresholds for the *emotion task* (*p < 0.01; *** p < 10^−6^; ns: no significant difference). KDEF images used in panels A and B: AF02NES, AF02HAS, AF02AFS.

These results indicate that individuals with MBS can reliably detect changes in facial features from static facial expressions (such as open or closed mouth) but are unable to efficiently process these changes to extract emotional information from them. Although static facial expressions are predominantly used in the literature to study emotion perception, in real life social interactions, we invariably encounter facial expressions in their dynamic form. Accordingly, we next sought to examine whether individuals with MBS who appear to have difficulty extracting emotion information from static faces, can accurately detect emotion from dynamic faces.

### Impaired perception of facial motion and facial emotion

Participants were shown video clips of happy, fearful, and angry faces (Figure 2A, 2B) of different durations in a randomized order, and asked to indicate if they thought the video clip depicted a happy, fearful, angry, or neutral expression (during the *emotion task* runs), or if they thought the mouth in the video moved or not (during the *motion control task* runs). The latter served as a perceptual control to establish whether individuals with MBS have an intact ability to perceive facial motion. We computed the average psychometric function for facial motion detection and facial emotion categorization (across all three emotions) for the perceptual control and emotion tasks, respectively, and then estimated the 50% threshold for detecting motion and categorizing emotion in the face. In addition, to examine any differences in emotion thresholds between expressions we fit the psychometric function for each expression and used this to estimate the 50% threshold for categorizing each emotion.

**Figure 2.**
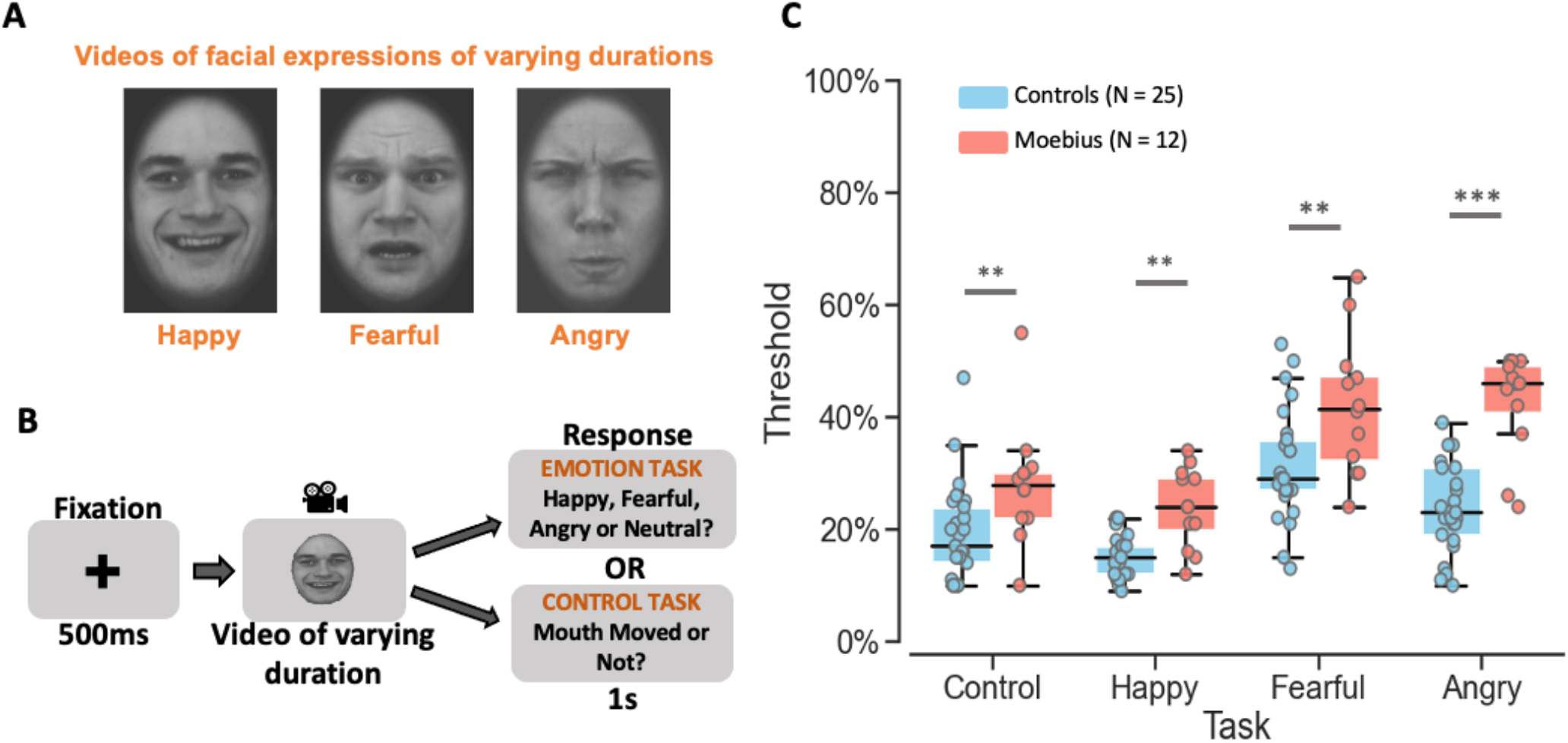
Dynamic Facial Expression Task. **A**. Videos of 8 different durations ranging from 200ms to 1.6s depicting dynamic facial expressions of happiness, fear, and anger were used as stimuli. **B**. Depiction of the trial structure – each trial began with a 500ms fixation cross, followed by a happy, fearful, or angry video of varying duration (200ms, 400ms, 600ms, 800ms, 1s, 1.2s, 1.4s, or 1.6s), and a 1s response window during which participants were to indicate if they thought the video depicted a happy, fearful, angry, or neutral expression during the *emotion task* runs, or whether the mouth moved or not during the *facial motion control task* runs. **C**. Box and whisker plots showing the thresholds for each task for control participants in blue and MBS individuals in red. MBS individuals had higher thresholds than controls participants for both facial *motion* detection and *emotion* categorization tasks (**p < 0.001; *** p < 10^−6^). KDEF images used in panels A and B: AM06HAS, AM17AFS, AF19ANS.

Overall, MBS individuals showed significantly higher thresholds than controls for both the emotion and perceptual control tasks, indicating that facial paralysis dampens the ability to extract both motion and emotion information from a face. A Task x Group mixed-effects beta regression analysis of facial motion and emotion thresholds (collapsed across expression type) showed not only an expected main effect of Task (Z = 2.81, p < 0.0049), but also a strong main effect of Group (Z = 4.67, p < 3.11 × 10^−6^), and no significant Task x Group interaction (Z = 0.60, p > 0.55), such that overall thresholds were significantly higher for MBS individuals compared to controls (50% thresholds for MBS: 31.2 ± 11.2% SD, Controls: 20.7 ± 7.7% SD), and significantly higher for the *emotion categorization task* compared to the facial *motion control task* (50% threshold for emotion detection: 28.3 ± 6.6% SD vs. mouth movement: 23.6 ± 10.1% SD). Further, within the *emotion task*, an Expression x Group mixed-effects beta regression analysis of categorization thresholds separated by expression type showed a main effect of Expression (p = 3.63×10^−16^), a main effect of Group (p = 1.59 × 10^−6^), and an Expression x Group interaction (p < 0.0038).

Post-hoc tests revealed that MBS individuals had significantly higher emotion categorization thresholds compared to controls for all three expression tasks, i.e., happy (t_79.9_ = 4.49, p < 2.37 × 10^−5^; 50% thresholds for MBS: 24.0 ± 6.9% SD vs. Controls: 15.3 ± 3.9% SD), fearful (t_79.9_ = 4.68, p < 1.15 × 10^−5^; MBS: 42.0 ± 12.3% SD vs. Controls: 31.3 ± 10.0% SD), and angry expressions (t_79.9_ = 8.48, p < 9.32 × 10^−^ _13_; MBS: 42.8 ± 9.1% SD vs. Controls: 24.0 ± 8.0% SD), with the latter yielding the largest difference in thresholds between the two groups. Critically, similar differences were seen in the control task as well, where post-hoc t-tests revealed higher facial motion detection thresholds for MBS individuals than the control group (t_57.5_ = 3.65, p < 0.00056; 50% thresholds for MBS: 28.0 ± 10.8% SD, Controls: 19.3 ± 8.7% SD). Thus, these results demonstrated that not only did MBS individuals have difficulty extracting emotion information from a moving face compared to controls, but they were also impaired at detecting facial motion from the dynamic face videos. These results indicate that facial paralysis in MBS impairs perception of facial motion in general, which in turn impacts perception of facial expressions. Next, we examined whether this deficit in detecting motion and recognizing emotion is limited to facial expressions, or whether it extends to other modes of social communication such as body expressions.

### Intact perception of body motion but not body emotion

In this task, participants were shown video clips of happy, fearful, and angry bodies of different durations (Figures 3A and 3B) in a randomized order and asked to indicate if they thought the actor’s body in the video clip depicted a happy, fearful, angry or neutral expression (during the *emotion categorization task* runs), or if they thought the actor’s arms in the video moved or not (during the *motion detection task* runs). The latter served as a perceptual control to establish whether individuals with MBS have an intact ability to perceive body motion. We computed the average psychometric function for body motion detection and body emotion categorization (across all three emotions) for the perceptual control and emotion tasks, respectively, and then estimated the 50% threshold for detecting motion and categorizing emotion in the body. In addition, to examine any differences in emotion thresholds between expressions we fit the psychometric function for each expression and used this to estimate the 50% threshold for categorizing each emotion.

**Figure 3.**
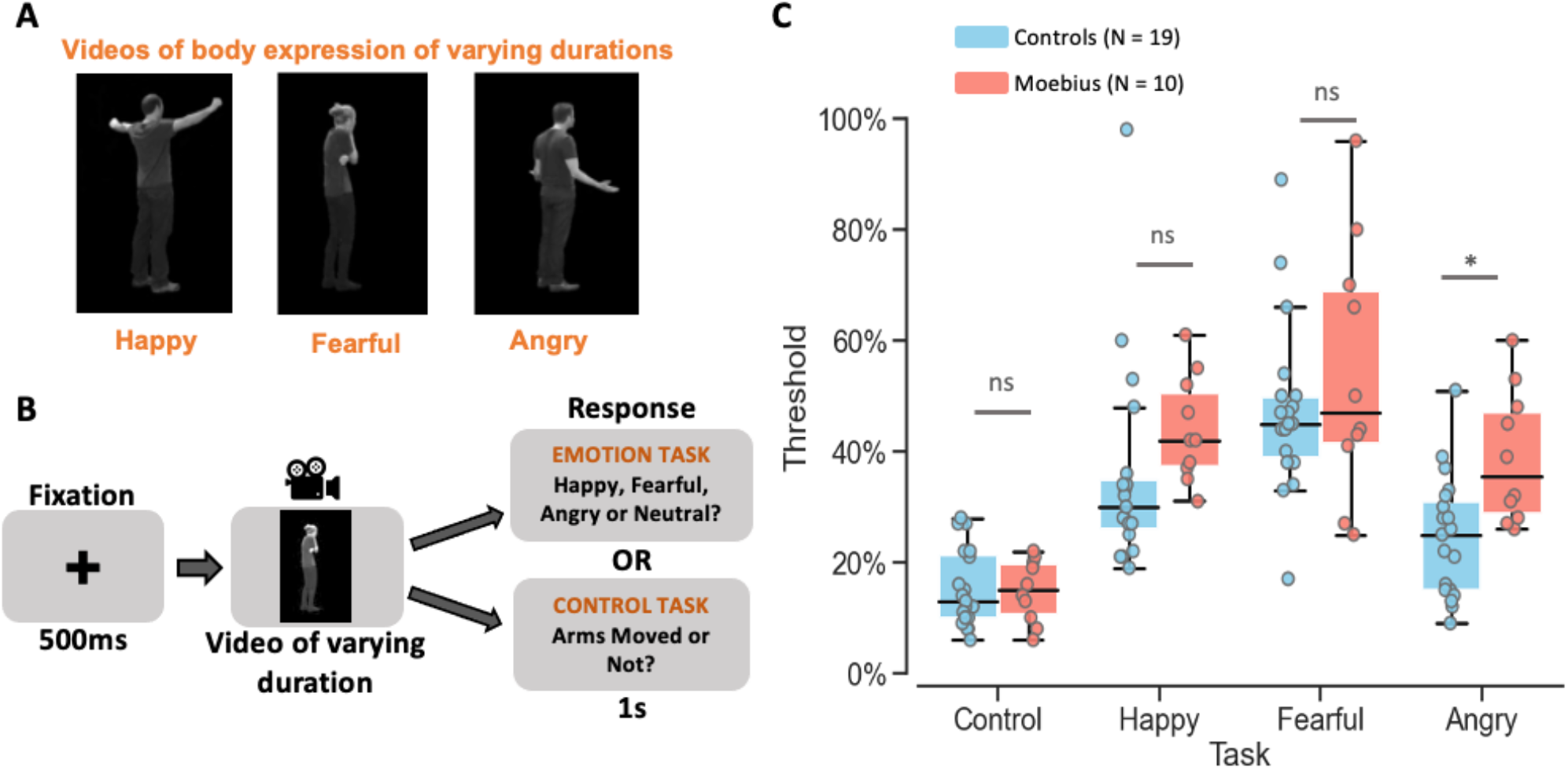
Dynamic Body Expression Task. **A**. Videos of 12 different durations ranging from 200ms to 2.4s depicting dynamic body expressions of happiness, fear, and anger were used as stimuli. **B**. Depiction of the trial structure – each trial began with a 500ms fixation cross, followed by a happy, fearful, or angry video of varying duration (200ms, 400ms, 600ms, 800ms, 1s, 1.2s, 1.4s, 1.6s, 1.8s, 2.0s, 2.2s, or 2.4s), and a 1s response window during which participants were to indicate if they thought the video depicted a happy, fearful, angry, or neutral expression during the *emotion task* runs, or whether the arms moved or not during the *body motion control task* runs. **C**. Graph showing the emotion thresholds for each task for control participants in blue and MBS individuals in red. MBS individuals had similar thresholds as control participants for *body motion* detection (*control task)* and for two of the three *body emotion* detection tasks (*p < 0.01; ns: no significant difference).

Overall, MBS individuals showed thresholds similar to controls for the *control task* but slightly higher thresholds for the *emotion* task, particularly for angry, indicating that although facial paralysis does not affect body motion perception, it dampens the ability to extract emotion information from body expressions. A Task x Group mixed-effects beta regression analysis of body motion and emotion thresholds (collapsed across expression type) showed the expected strong main effect of Task (Z = 10.26, p < 2.0 × 10^−16^), no main effect of Group (Z = 1.33, p > 0.18), and no significant Task x Group interaction (Z = 1.70, p < 0.09), such that detection thresholds were significantly higher for the *body emotion task* compared to the *body motion control task* (50% thresholds for emotion detection: 39.6 ± 11.4% SD, arm movement: 15.1 ± 6.98% SD). While the interaction effect did not reach statistical significance, there was a trend for MBS individuals to have numerically higher thresholds than controls on the *emotion task* but not the *motion task*. Further, within the *emotion categorization task*, an Expression x Group mixed-effects beta regression analysis of thresholds separated by expression type, showed a main effect of Expression (p < 8.33 × 10^−6^), a main effect of Group (p < 0.04), and no Expression x Group interaction (p > 0.24). Thus, these results showed that although MBS individuals did not have difficulty extracting *motion* information from a moving body, they suggest that they may have some trouble extracting *emotional* cues from the body motion information.

These results, combined with results from the *static* and *dynamic facial expression* tasks, indicate that facial paralysis in MBS selectively impairs perception of facial motion and does not impact body motion perception. However, the inability to portray emotion on their own face dampens the ability of MBS individuals to extract emotional information from both facial and body expressions.

Having demonstrated a strong impairment in detecting and recognizing emotion from facial and body expressions, we next sought to probe whether these deficits were limited to emotion detection and categorization tasks, or whether the impairment transcends into other types of more complex tasks such as expression matching. In addition, since all three experiments employed simple feature and motion detection as controls tasks, in the next experiment we used the more complex visual process of facial identity matching as a control task. This served to identify any generalized high-level processing deficits in MBS individuals relative to controls.

### Selective impairment in higher order face processing: facial identity vs. facial expression

In this task, participants were shown three faces in a triangular formation (Figure 4A and 4B) and asked to indicate which of the two lower faces (or neither) matched the top face based on the facial identity of the actor (during *identity task* runs) or based on facial expression of the actor (during *expression task* runs). Overall, MBS individuals performed similar to controls on the *identity matching* but worse on *expression matching* (Figure 4C). A Task x Group mixed-effects beta regression analysis of accuracy rates (percent correct) showed a main effect of Task (Z = 4.26, p < 2.05 × 10^−5^), no main effect of Group (Z = 1.02, p > 0.31), and a significant Task x Group interaction (Z = 2.14, p < 0.03). Post-hoc between group comparisons for each of the tasks showed that MBS individuals (mean ± SD: 60.2 ± 17.3%) performed similar to controls (mean ± SD: 60.5 ± 18.6%) on the *identity task* (t_24.3_ = 0.10, p > 0.93), but were much worse on the *expression task* (MBS: 67.5 ± 8.3% SD; controls: 79.1 ± 8.1% SD; t_24.3_ = 2.998, p < 0.006). These data indicate that individuals with MBS have difficulty processing emotional cues and suggest that other higher order face processing functions such as identity processing are not impacted by the facial paralysis.

**Figure 4.**
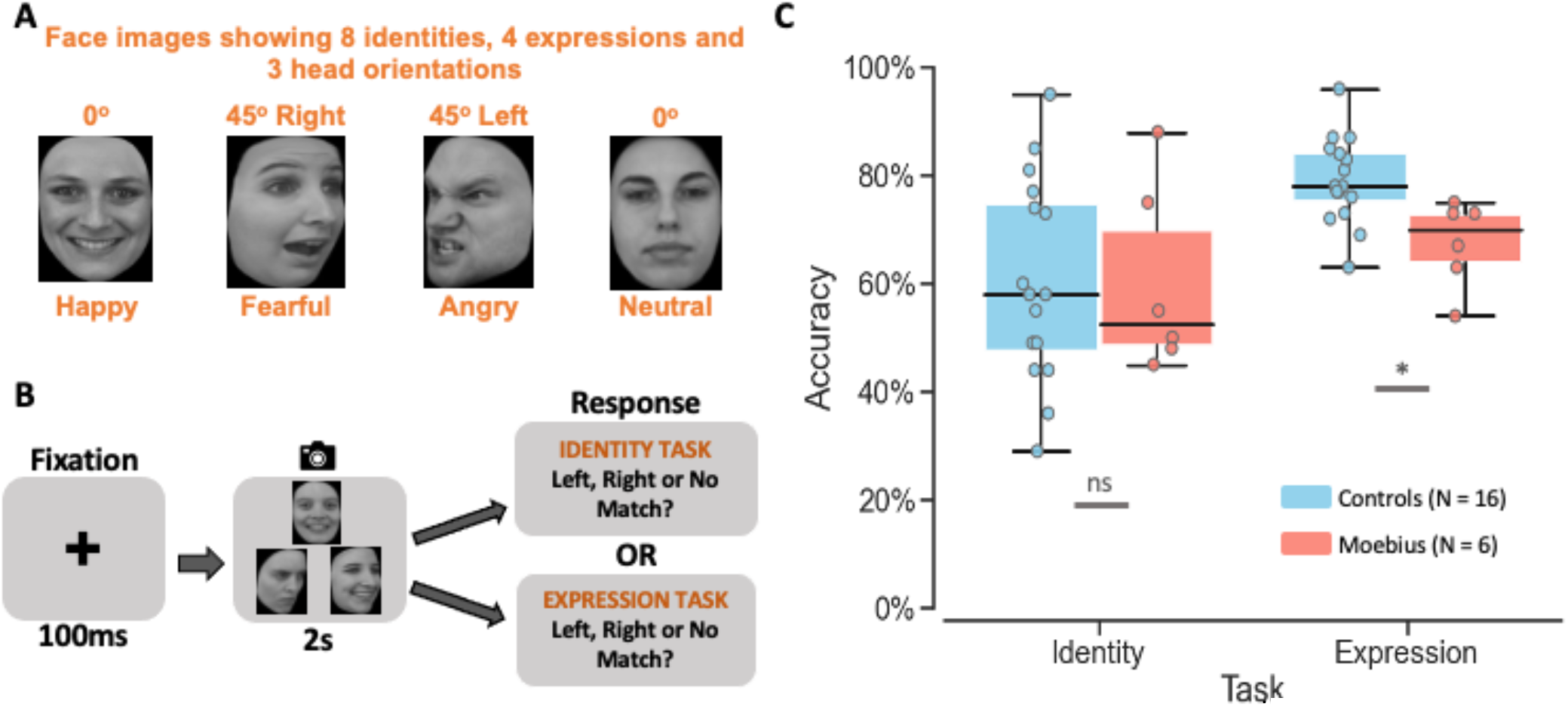
Identity and Expression Matching Task. **A**. Images of 8 different facial identities and 4 different expressions (happy, fearful, angry, and neutral) from 3 different viewpoints (left, front, and right) were used as stimuli. **B**. Depiction of the trial structure – each trial began with a 100ms fixation cross, followed by the presentation of three faces in a triangle format for 2s, during which participants were to indicate which of the two bottom faces (or neither) matched the top face either on *identity* or *expression* (in separate runs). **C**. Box and whisker plots showing performance accuracy for *identity* and *expression* matching for control participants in blue and MBS individuals in red. MBS individuals had similar performance as control participants on the *identity matching task* but worse performance on the *expression matching task* (*p < 0.01; ns: no significant difference). KDEF images used in panels A and B: AF01HAS, AF13AFHR, AM17ANHL, AF17NES, AF21HAS, AF21ANHR, AF13HAHR.

Overall, the results from our comprehensive battery of behavioral tasks suggested that facial paralysis (as seen in MBS) results in a dampening of emotion processing, which is evident from higher thresholds on emotion detection and recognition tasks and from reduced performance on an expression matching task. But what brain changes, if any, subserve this generalized dampening in emotion processing in MBS? To answer this question, we conducted exploratory fMRI scans to examine the potential neural correlates of this behavioral impairment in a small subset of MBS individuals and age- and gender-matched controls, focusing on face-selective regions of the brain^23-25^ such as fusiform face area (FFA), posterior superior temporal sulcus (pSTS) and the amygdala (AMG).

### Potentially reduced amygdala engagement during facial expression processing

Using a functional face localizer, we first identified face-selective regions of interest (ROIs) in the right FFA and right pSTS that responded significantly more to static and dynamic faces than objects (see *Methods* for details) for each participant. In addition, using a standard neuroanatomical atlas (*TT_N27* in AFNI), we defined an anatomical ROI in the right AMG for each participant. Within these three ROIs, we then compared the average fMRI activity during *identity* vs. *expression tasks* for controls and individuals with MBS.

Overall, the right FFA and right pSTS showed similar fMRI activity profiles for the two tasks between controls and MBS, while activity profiles in the amygdala appeared different between the two groups. A Task x Group linear-mixed effects ANOVA of right FFA data showed no significant main effect of Task (p > 0.77) or Group (p > 0.97), or Task x Group interaction (p > 0.22). fMRI activity in right pSTS expectedly showed a significant main effect of Task (p < 0.01), but no main effect of Group (p > 0.97) or Task x Group interaction (p > 0.89). Due to the small sample size for the MBS group, we also compared the fMRI activity within each group between the two tasks. These paired t-tests revealed no difference in activity between *Identity* and *Expression* conditions in right FFA for both controls (p > 0.23) and MBS (p > 0.23). By contrast, in right pSTS, controls showed higher fMRI activity during *expression* than *identity task* runs, while in the MBS group this comparison did not reach statistical significance (Controls: t_14_ = 2.46, p < 0.01; MBS: t_5_ = 1.87, p < 0.06; see Figure 5A and 5B). In contrast to right FFA and right pSTS, a different pattern was observed in right AMG. While no significant main effect of Task (p > 0.78), main effect of Group (p > 0.49) or Task x Group interaction (p < 0.07) was found in right AMG, the two groups showed seemingly opposite fMRI activity profiles. Although these effects were not statistically significant, in contrast to control participants who on the average showed numerically higher activity during *expression* than *identity task* runs (10 out of 15 controls showed this pattern; paired sampled t-test: t_14_ = 1.41, p < 0.09), individuals with MBS predominantly showed the opposite pattern, i.e., numerically lower activity during *expression task* runs than during *identity task* runs (4 out of 6 MBS individuals showed this opposite pattern; paired sample t-test: t_5_ = −1.81, p < 0.07; see Figure 5C). Due to the low number of MBS participants, we also ran non-parametric Wilcoxon signed rank tests for the right AMG and found similar results (controls: Z = 1.48, p < 0.07; MBS: Z= −1.57, p < 0.06). In addition, we also conducted Bayesian prevalence analysis^26^ and found no significant prevalence difference between control and MBS groups for any of the regions (see Supplemental Material E). Here too however, we found a qualitatively different pattern of results in the right AMG for MBS individuals, who showed zero prevalence of the expected *expression* > *identity* effect compared to controls who showed positive prevalence. Thus, taken together, although the fMRI results are inconclusive and should be interpreted with caution due to low statistical power, these preliminary data suggest that while MBS individuals show relatively normal activation patterns in face-selective regions such as FFA and pSTS, the amygdala which is known to be a core region for processing emotion in general and facial expressions in particular, may exhibit a different pattern of activation compared to controls.

**Figure 5.**
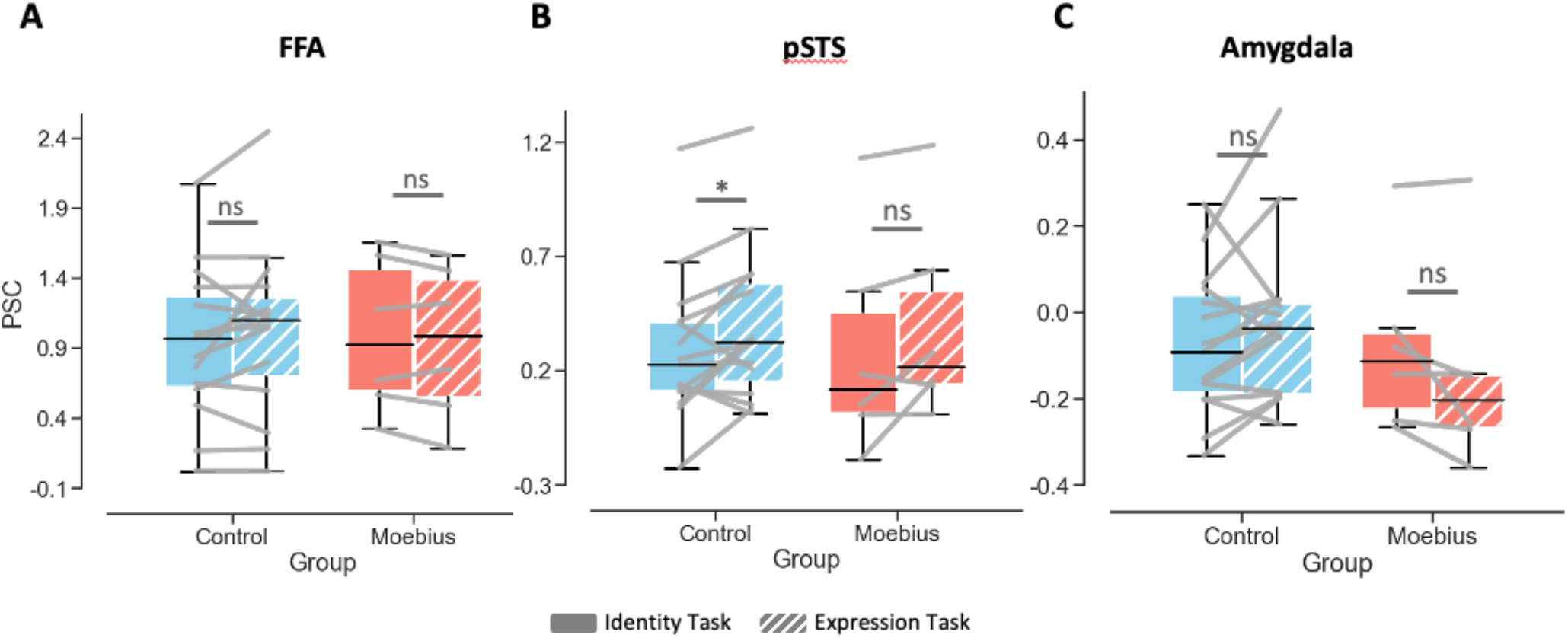
Neural correlates of expression processing in a small group of MBS individuals relative to controls. **A**. fMRI activity plots showing no difference in BOLD percent signal change in right FFA during *identity* (solid) vs. *expression* (patterned) *matching* for Controls (N = 15) in blue and MBS individuals (N = 6) in red. **B**. fMRI activity plots showing significantly higher BOLD percent signal change in right pSTS for *expression* compared to *identity matching* for Controls (*p < 0.01) but not MBS (p = 0.06). **C**. BOLD percent signal change in right amygdala was not significantly different during *expression* matching relative to *identity* matching for Controls or MBS, although numerically the two groups showed opposite patterns (Controls: *expression* > *identity* while MBS: *expression* < *identity*).

## DISCUSSION

Using a battery of behavioral tasks, we found that individuals with Moebius Syndrome, who from birth are unable to move their facial muscles to express emotions, were impaired at detecting emotion in others’ facial expressions. This impairment in facial emotion detection stemmed from an impairment in facial motion perception. Importantly, while the facial paralysis did not affect the ability of MBS individuals to perceive body motion, it did impact their ability to extract emotional information from body expressions. Thus, the inability to move a particular body part (in this case one’s face) impacts the perception of motion and emotional information in the same body part of others (here facial expressions), and further, may result in a generalized dampening of emotion perception (demonstrated here with body expressions). Additionally, our MBS participants although impaired at facial expression processing, could accurately perform other challenging perceptual tasks such as feature detection and identity matching. These behavioral findings were supported by preliminary fMRI data in a small subgroup of MBS individuals that suggested reduced engagement of the amygdala during an expression processing task. Taken together, these convergent findings suggest that the inability to make facial expressions from birth, results in perceptual dampening of others’ facial expressions and is associated with changes in the level of engagement of the neural circuitry underlying emotion processing.

Our results of dampening of emotion perception align well with those of others who have previously reported generalized face processing difficulties in Moebius Syndrome^20^ as well as those reporting reduced autonomic modulation during emotion processing^21,22,27^. Our results are, however, at odds with the conclusion of some studies that have reported no impact of facial paralysis on emotion perception in Moebius Syndrome^16-18^. However, the use of full-blown facial expression stimuli, lack of appropriate control tasks and small sample size in these previous studies may account, in part, for their negative results. Vannuscorps et al.,^18^ did limit the amount of emotion information in one of their facial expression tasks (40% of full-blown facial expressions), but our data suggest that this may not have been low enough to pick up the subtle emotion deficit in MBS. In our study, most MBS individuals were, on average, able to perform the task at 40% morph level (even though on the average their thresholds were higher than controls). Thus, the task used by Vannuscorps et al.^18^ likely was not sensitive enough to pick up subtle deficits in emotion processing. Individuals with MBS do not typically report significant difficulties in social interactions^17^, and thus it is not surprising that their emotion perception deficits are subtle and that they are able to efficiently recognize full blown emotional expressions. Affected individuals have likely developed compensatory strategies during development to not only convey their own emotions, but also to understand another person’s emotional state by using other social cues such as voice tonality^28^. Although we did not collect physiological measures of emotion perception in the current study, future studies using autonomic skin conductance (as a proxy for emotional reactivity), eye tracking and electromyographic data, will help to further explicate the link between emotion perception and emotional experience. Future work could also focus on the generalizability of the dampening that we observed in the visual domain in our current study by examining the effect of facial paralysis on emotion perception in other modalities, such as perception of emotional vocalizations.

Further, with regard to the choice of task type for our questions of interest, we used a yes-no detection task for static faces, and a 4-way emotion categorization task for dynamic faces and bodies; tasks that are susceptible to user bias. Typically, a two-interval forced choice (2-IFC) task is often recommended for such psychophysical experiments due to its limited susceptibility to response bias^29,30^ (but also see Yeshurun et al.^31^ for why 2-IFC procedures may not be completely bias-free). However, we chose our tasks in this way for their simplicity and short administration times, and so that we could limit task demand and cognitive load confounds. We also confirmed that the bias inherent in the task was not significantly different between the two groups by performing signal detection theory analysis on the catch trials in the static face task (see Supplemental Material C). Importantly, we also found that the sensitivity (d’) to emotional faces did not vary between the two groups, which further supports the notion that the emotion deficits in MBS are subtle, and that individuals do not experience difficulty in detecting emotion from full-blown static facial expressions as has been used in several prior studies. While we were able to show that response bias could not explain the differences between the two groups for the static face task, a similar analysis for the dynamic face and body tasks was not possible due to the absence of neutral videos in these tasks. Thus, although unlikely, it is possible that bias could have played a role in the results we observed for the dynamic tasks. Another potential limitation of the static-face task was the estimation of thresholds based on psychometric fitting of staircase data which results in sub-optimal sampling of data along the psychometric range. This issue was alleviated by the use of the method of constant stimuli in the dynamic face and body tasks.

Overall, our results also align well with the general notion of experience influencing perception as has been demonstrated by studies in professional actors^13^ and dancers^14,15^ showing enhanced ability to recognize facial and body expressions, respectively. Results from these studies on how experience enhances perception therefore predict that the lack of facial experience in MBS would adversely impact the ability to recognize others’ facial expressions. In fact, these results provide direct support for Darwin’s intuitive observation^32^ that suppressing one’s expressions of an emotion also suppresses the underlying emotion (facial feedback). Darwin also held that people tend to expressively imitate others (facial mimicry), and the current study in individuals with congenital facial palsy, who have never been able to imitate or express their emotions, finds what Darwin had predicted – a dampened sensitivity to facial emotional information in such individuals. Our results are in line with a recent study describing the role of motor control in visual body perception that reported impaired performance on a body judgement task in a group of congenital one-handers^33^. In fact, the crucial role of motor experience in emotion processing has been clinically leveraged in the treatment of depression by using muscle inactivation to manipulate emotional proprioception^34^.

In contrast to previous studies of emotion processing in Moebius Syndrome, our study has the added value of neuroimaging to examine the neural correlates underlying the impact of facial paralysis on emotion perception. Although the fMRI data are limited in terms of sample size (only 6 MBS participant completed the *identity* and *expression tasks* in the scanner), they suggest that a more robust investigation of the amygdala and related emotion processing circuitry in MBS is warranted in the future. Additionally, the trend for the *expression* vs. *identity* effect in pSTS to be slightly weaker in MBS than controls, should also be further studied as it may be linked to the potentially diminished involvement of the amygdala in MBS and reflect their behavioral deficit. Since it appears that the degree of engagement of these regions (especially the amygdala) may be impacted in MBS relative to controls, future studies could examine the use of visual training to improve the engagement of these networks during emotion processing.

In summary, our behavioral and neuroimaging findings together provide support for a critical role of facial movement (and relatedly facial mimicry, motor experience, and facial feedback) in facial motion perception such that its disruption can lead to impairment in the perception of facial expressions. These results predict the effectiveness of visual training paradigms to enhance emotion perception and improve social interactions in situations involving absent motoric experiences, such as the congenital facial palsy seen in Moebius Syndrome.

## MATERIALS AND METHODS

### Participants

Twenty-four individuals with Moebius Syndrome (17-64 years) and 50 normal volunteers (20-63 years) (or a legal guardian where applicable) provided written informed to participate in the study. MBS participants were enrolled under a protocol approved by the Institutional Review Board of the National Human Genome Research Institute at the National Institutes of Health (NCT02055248). MBS individuals (see supplemental material for clinical information) were recruited to the study at the NIH Clinical Center (N=14) or at the 2018 MBS Foundation Conference in St. Petersburg, FL (N=10). Some of the MBS participants (N=16) completed survey questionnaires about their medical history and facial disability and some (N=10) completed additional neuropsychological testing. Healthy volunteers for the study were recruited under an NIH IRB approved protocol of the National Institute of Mental Health (NCT00001360). MBS and control participants completed a series of computer tasks involving static and dynamic facial and body expressions. Due to time and availability constraints, not all participants completed all tasks. Experimental conditions were kept similar between the two testing venues (NIH Clinical Center and MBS Foundation Conference). Eight MBS individuals and 16 control participants also completed functional magnetic resonance imaging (fMRI) scans at the NIH Clinical Center.

### Static facial expression task

Fourteen MBS participants (mean age 35.2 ± 17.7 SD, 6 males) and 26 age- and gender-matched controls (mean age 32.1 ± 13.3 SD, 10 males) completed 2 runs of an *emotion detection task* and 2 runs of a *feature detection task* (that served as a perceptual control task) shown in Figure 1. Using FantaMorph software (Abrosoft) we first created a series of morphs with varying levels of happiness and fear using a neutral face and the corresponding happy or fearful face, respectively. Neutral and full-blown facial expression images for 14 male and 13 female actors were taken from the Karolinska Directed Emotional Faces set (KDEF^35^). For each actor, 19 intervening morphs of increasing expressional content in 5% increments were created by morphing the neutral image to the corresponding happy and fearful images. Starting at the 50% morph, participants were shown images of increasing or decreasing emotion depending on their performance in a 3-down/1-up adaptive staircase fashion. Each run contained a total of 169 trials consisting of 130 morph face trials (dependent on participant’s performance), 26 neutral face catch trials, and 13 full expression catch trials. Each trial began with a fixation cross for 250ms followed by the morphed face. The face appeared at the center of the screen for 350ms and subtended 4 and 6 degrees of visual angle in the horizontal and vertical dimensions, respectively. The face was followed by a fixation cross for 1.5s during which participants were asked to press the “yes” button if they thought the image showed a happy or fearful expression (in separate runs) or press the “no” button if they thought the image was neutral (see Figure 1A and 1B for stimuli and task details). In separate controls runs, using a similar procedure, participants were asked to press the “yes” button if they thought the image showed an open mouth (lips apart) or press the “no” button if they thought the image showed a closed mouth (lips touching). Responses were recorded using a response box (Current Designs, Inc.) and participants were asked to respond as quickly and as accurately as possible. Detection accuracies for each morph level for fearful and happy faces were used to generate each participant’s psychometric curves. A standard logistic function was used to fit each participant’s psychometric curve for each emotion using the *fminsearch* function available in the MATLAB Optimization Toolbox (MathWorks, Inc.). The α parameter of this fit (which denotes the point of subjective equality for a yes-no detection task^36,37^) was then used to identify the morph level threshold at which each participant achieved 50% detection accuracy. One MBS participant’s data were excluded from further analysis because the fitting procedure did not converge to yield appropriate parameter estimates. Morph level thresholds for the remaining participants were then entered into a mixed-effects beta regression model (using the R package *mgcv*) with Task and Expression as within-subjects factors and Group as a between-subjects factor (see results in Figure 1C). Additionally, where appropriate, post-hoc Bonferroni-corrected pairwise comparisons were conducted to compare average thresholds between Controls and MBS groups for each emotion.

### Dynamic facial expression task

Thirteen MBS participants (mean age 33.5 ± 16.5 SD, 6 males) and 25 age- and gender-matched controls (mean age 31.9 ± 11.8 SD, 11 males) completed 3 runs of a *facial emotion task* and 1 run of a *facial motion control task* shown in Figure 2 (one MBS participant completed only 2 runs of the *emotion task* due to time constraints). Stimuli for this task were created by stringing together the 21 morph images for each expression (created for the static facial expression task described above) into a slow-motion movie depicting the actor making a happy, fearful, or angry facial expression. The movie was then clipped to create 8 different lengths of the expression, such that each clip showed either 200ms, 400ms, 600ms, 800ms, 1s, 1.2s, 1.4s and 1.6s from the start of the movie. Shorter clips contained less emotion information than longer clips since the expression on the face in the video had not yet evolved into the full-blown expression. In this way, we limited the amount of emotion information in the dynamic facial expression movies. Using a method of constant stimuli, videos of varying lengths of the three different expressions were presented to participants in a randomly inter-mixed order within each run. Each trial began with a 500ms fixation cross, followed by the video clip presented at the center of the screen subtending 4 (horizontal) and 6 (vertical) degrees of visual angle. The video was in turn followed by a response window lasting 1s, during which participants were asked to indicate their response (see Figure 2A and 2B for stimuli and task details). Each run contained 6 trials of each video length of each emotion, totaling 144 trials per run. During emotion categorization task runs, participants were asked to press one of four buttons to indicate whether the video clip depicted a happy, fearful, angry, or neutral expression. During facial motion detection task runs participants were asked to press one of two buttons to indicate whether the actor’s mouth moved during the video clip. Responses were recorded using a four-button response box (Current Designs Inc.) and participants were asked to respond as quickly and as accurately as possible. Data were analyzed similar to the *static facial expression task* such that detection and categorization accuracies for each video duration for each emotion were first used to generate each participant’s psychometric curves. The α parameter of a standard logistic function fit to the psychometric curves was then used to identify the video duration threshold at which each participant achieved 50% accuracy for each emotion. One MBS participant’s data were excluded from further analysis because the fitting procedure did not converge to yield appropriate parameter estimates. Duration thresholds for the *facial motion detection* and *facial emotion categorization tasks* were entered into a mixed-effects beta regression analysis with Task (2 levels: control task and emotion task) as a within-subjects factor and Group as a between-subjects factor. Further, within the emotion categorization task, thresholds for the three different expression types (happy, fearful, and angry) were entered into a 3 Expression x 2 Group mixed-effects beta regression analysis. Additionally, where appropriate, Bonferroni-corrected post-hoc t-tests were conducted to compare average thresholds between Controls and MBS groups for each task condition (see results in Figure 2C).

### Dynamic body expression task

Eleven MBS participants (mean age 37.7 ± 16.0 SD, 4 males) and 20 age- and gender-matched controls (mean age 35.1 ± 12.8 SD, 6 males) completed 3 runs of a *body emotion task* and 1 run of a *body motion control task* shown in Figure 3 (one MBS participant completed only 2 runs of the *emotion task* due to time constraints). Stimuli for this task were chosen from the *Action Database* of whole-body action videos^38^ of 10 actors making happy, fearful, and angry body movements (with their backs towards the camera to obscure the face). Each video was then clipped to create 12 different lengths of the body expression, such that each clip showed either 200ms, 400ms, 600ms, 800ms, 1s, 1.2s, 1.4s, 1.6s. 1.8s, 2s, 2.2s or 2.4s from the start of the video. Shorter clips contained less emotion information than longer clips since the movement of the body in the video had not yet evolved into a full-blown expression. In this way we limited the amount of emotion information in the dynamic body expression videos. Using a method of constant stimuli, videos of varying lengths of the three different expressions were presented to participants in a randomly inter-mixed order within each run. Each trial began with a 500ms fixation cross, followed by the video clip presented at the center of the screen subtending 6 (horizontal) and 4 (vertical) degrees of visual angle. The video was in turn followed by a response window lasting 1s, during which participants were asked to indicate their response (see Figure 3A and 3B for stimuli and task details). Participants completed 10 trials for each video length for each emotion, over 3 runs of the emotion task, with each run containing 120 trials. During emotion categorization task runs, participants were asked to press one of four buttons to indicate whether the video clip depicted a happy, fearful, angry, or neutral expression. During facial motion detection task runs participants were asked to press one of two buttons to indicate whether the actor’s mouth moved during the video clip. Responses were recorded using a four-button response box (Current Designs Inc.) and participants were asked to respond as quickly and as accurately as possible. Data were analyzed similar to the *dynamic facial expression task* such that categorization accuracies for each video duration for each emotion were first used to generate each participant’s psychometric curves. The α parameter of a standard logistic function fit to the psychometric curves was then used to identify the video duration threshold at which each participant achieved 50% accuracy for each emotion. One MBS and one Control participant’s data were excluded from further analysis because the fitting procedure did not converge to yield appropriate parameter estimates. Duration thresholds for the *body motion detection* and *body emotion categorization tasks* were entered into a mixed-effects beta regression analysis with Task (2 levels: control task and emotion task) as a within-subjects factor and Group as a between-subjects factor. Further, within the emotion categorization task, thresholds for the three different expression types (happy, fearful, and angry) were entered into a 3 Expression x 2 Group mixed-effects beta regression analysis. Additionally, where appropriate, Bonferroni-corrected post-hoc t-tests were conducted to compare average thresholds between Controls and MBS groups for each task condition (see results in Figure 3C).

### Facial identity and expression matching task

Six MBS participants (mean age 46.2 ± 18.1 SD, 3 males) and 16 age- and gender-matched controls (mean age 38.1 ± 14.3 SD, 8 males) completed 2 runs each of a *facial identity matching task* and a *facial expression matching task* shown in Figure 4. Stimuli for this task were prepared by selecting images of 8 actors depicting happy, fearful, angry, and neutral facial expressions from 3 viewpoints (front view and 45° to left and right) from the KDEF database. On each trial, participants were shown 3 faces in a triangular formation for 2s (preceded and followed by a 100ms fixation cross; see Figure 4A and 4B) and were asked to indicate which of the two lower faces (or neither) matched the top face based on the facial identity of the actor (during *identity task* runs) or based on facial expression of the actor (during *expression task* runs). The top face was one of the 8 actors depicting one of the 4 expressions (from one of the 3 viewpoints), and one of the bottom faces matched the top face on identity (same actor, different expression) and the other matched the top on expression (different actor, same expression). On some trials neither of the bottom faces matched the top on identity or expression, and these served as catch trials to prevent participants from performing this task without explicitly detecting identity or expression. Each face in the triangular display subtended 2 (horizontal) and 3 (vertical) degrees of visual angle at 3-degree eccentricity about the central fixation cross. Each run consisted of 8 blocks of 12 trials, with each trial lasting 2.2s. Group differences in performance (accuracy rates) on each task were assessed using a mixed-effects beta regression analysis with Task (2 levels: identity and expression) as a within-subjects factor and Group as a between-subjects factor, as well as post-hoc Bonferroni-corrected pairwise comparisons.

### Functional MRI data acquisition

Eight individuals with MBS (mean age 37.1 ± 22.2 SD, 4 males) and 16 age- and gender-matched controls (mean age 32.9 ± 14.1 SD, 8 males) were scanned using a GE-MR750 3 Tesla MRI scanner. While in the MRI scanner, participants completed two blocked-design functional face localizer task runs, and 2 runs each of the *identity* and *expression matching* task (one MBS individual did not complete this task due to time constraints). fMRI scans were performed using a BOLD-contrast sensitive multi-echo echo-planar (EPI) sequence with three echo times (TEs: 12.5ms, 27.6ms and 42.7ms). Scanning parameters used were typical of whole brain fMRI studies (Array Spatial Sensitivity Encoding Technique [ASSET] acceleration factor = 2; TR = 2.2s; 33 interleaved AC-PC aligned 3.5mm thick slices with 3.2 × 3.2mm in-plane resolution). One participant (Patient 1) was scanned using a single echo EPI sequence (TE: 27.6ms) due to unavailability of the multi-echo sequence at the time, but all other parameters remained the same. A high-resolution T1 structural MP-RAGE scan (172 sagittal slices with 1mm x 1mm x 1mm voxel resolution, TE = 3.47ms, TR = 2.53ms, TI = 900ms, flip angle = 7°) was also collected for each participant.

During each *Face localizer* run, participants were shown in a random order, 2 blocks each of videos of moving faces, moving objects, static faces, static objects, or scrambled images. Each block within the run lasted 26.4s and contained 12 stimuli presented for 2s each with 200ms fixation between stimuli. Between blocks of stimuli participants were asked to fixate during baseline blocks lasting 13.2s each, resulting in each run lasting about 7min. The stimuli appeared at the center of the screen subtending 6 and 8 degrees of visual angle in the horizontal and vertical dimensions, respectively. Participants were instructed to perform a simple one-back matching task during the stimulus blocks and indicate with a button press whether the current image was the same as the previous image, or not. Participants also completed two each of the *identity* and *expression matching* tasks (presented in an alternating fashion) similar to the task outside the scanner (see Figure 4A and 4B). Each run consisted of 8 blocks of 12 trials each that lasted 28.8s (2.2s each for each trial within a block), followed by 13.2s of intervening baseline fixation, resulting in each run lasting about 6min 48s. Prior to the experiment participants practiced the tasks outside the scanner and were instructed to respond as quickly and accurately as possible while fixating at the central fixation cross.

### Functional MRI data analysis

Each participant ‘s fMRI data were preprocessed and analyzed using multiple programs in AFNI (Analysis of Functional Neuroimages^39,40^). Preprocessing of fMRI data was performed using methods similar to those previously reported using AFNI’s *afni_proc*.*py* program^41^. In short, fMRI data were first despiked, the first four volumes of each run were discarded, and the remaining volumes were distortion corrected, slice-timing corrected, and registered to each other. Structural and functional data were first aligned and then spatially normalized to a Talairach-space aligned version of the *ICBM_152_2009c* atlas template^42^ using a non-linear warping procedure. For the task runs, data for the three echoes were optimally combined using standard methods^43^, then smoothed using a 4-mm FWHM smoothing kernel. The resulting time series were then normalized by the mean signal intensity of each voxel to reflect percent signal change, which served as the input for subsequent regression analysis using a General Linear Model (GLM).

### Amygdala and face-selective ROI definition

Data from both localizer runs were concatenated into one GLM analysis which modeled five conditions of interest (*Dynamic faces, Dynamic objects, Static faces, Static objects*, and *Scrambled images*) using standard hemodynamic response functions of 26.4s duration, 3 baseline parameters (second-order polynomial) per run, and six motion regressors of no-interest per run. The localizer GLM results were used to functionally define face-selective regions of interest (ROIs) in right fusiform face area (rFFA) and right posterior superior temporal sulcus (rpSTS) for each participant, by first thresholding the statistical maps of fMRI activity of faces relative to objects (t-value of contrast between dynamic and static faces vs. dynamic and static objects) using an individual voxel threshold of p < 0.001 and a cluster-corrected α of 0.05 (resulting in cluster size thresholds of 11 to 20 voxels per participant). Then, to locate right FFA and pSTS, activation clusters in the fusiform gyrus and posterior portions of the superior temporal sulcus, respectively, were identified, and the location of their peak voxels noted. For right amygdala, a standard Talairach atlas^44^ (*TT_N27*) was used to anatomically define a ROI in right amygdala (AMG).

### Identity vs. expression task fMRI analysis

Data from the 4 identity and expression matching runs were concatenated into one GLM analysis which modeled 2 conditions of interest (Identity and Expression) using standard hemodynamic response functions of 28.8s duration, 3 baseline parameters (second-order polynomial) per run, and six motion regressors of no-interest per run. Data for one control participant and one MBS individual were excluded from further analysis because of significant loss of degrees of freedom due to movement-and noise-related censoring of timepoints. Using the anatomically defined amygdala ROI and a 4mm radius sphere drawn around the peak activated voxel in the right FFA and right pSTS face-selective ROIs, we extracted out the mean fMRI percent signal change (beta coefficients from GLM analysis) within these regions, during the *Identity* and *Expression* task conditions, separately. We performed a linear mixed-effects Task x Group analysis and paired t-tests within each group of participants to determine whether these regions were engaged to the same degree during *identity* and *expression* processing.

## ACKNOWLEDGEMENTS

This research was supported by the NIH Division of Intramural Research under NHGRI protocol 14-HG-0055 (ZIA# HG200389; NCT02055248) and NIMH protocol 93-M-0170 (ZIA# MH002918; NCT00001360). The Moebius Syndrome Research Consortium (members detailed in supplemental material) received research funding from the National Institutes of Health (Grant Number: U01HD079068). We thank Dr. Kathleen Rives Bogart for granting us use of the Moebius Syndrome survey, as well as for thought provoking discussions. Many thanks to Flavia Facio, Carol Van Ryzin, and Jose Salas for their help in recruiting and scheduling participants, and to the Moebius Syndrome Foundation for their assistance and support. We are grateful to all the Moebius Syndrome individuals and their families for their generous participation. Thanks to Shivani Goyal and Hannah Wild for their help with survey data compilation.

## AUTHOR CONTRIBUTIONS

SJ, CIB, and LGU conceived and designed the study; SJ, JJ, JL, and SL performed the experiments and collected data; GC provided statistical advice; EWJ, BWD, ECE, IM, JS, and MSRC recruited and characterized individuals with Moebius Syndrome; SJ, and JJ, analyzed data; SJ, wrote the paper; all authors edited the paper; IM and LGU secured funding.

## DATA AND CODE AVAILABILITY

All data reported in this paper will be shared by the lead contact upon request. Original analysis code is available from the lead contact upon request. Any additional information required to reanalyze the data reported in this paper is available from the lead author upon request.

## BIBLIOGRAPHY

1. Soderling, B. (1959). The first smile; a developmental study. Acta Paediatr Suppl 48, 78–82.

2. Tautermannova, M. (1973). Smiling in infants. Child Dev 44, 701–704.

3. Wolff, P. (1963). Observations on the early development of smiling.. In Determinants of infant behavior.

4. Dimberg, U., and Thunberg, M. (1998). Rapid facial reactions to emotional facial expressions. Scand J Psychol 39, 39–45. 10.1111/1467-9450.00054.

5. Gallese, V., and Sinigaglia, C. (2011). What is so special about embodied simulation? Trends Cogn Sci 15, 512–519. 10.1016/j.tics.2011.09.003.

6. Ross, P., and Atkinson, A.P. (2020). Expanding Simulation Models of Emotional Understanding: The Case for Different Modalities, Body-State Simulation Prominence, and Developmental Trajectories. Front Psychol 11, 309. 10.3389/fpsyg.2020.00309.

7. Keysers, C., and Gazzola, V. (2009). Expanding the mirror: vicarious activity for actions, emotions, and sensations. Curr Opin Neurobiol 19, 666–671. 10.1016/j.conb.2009.10.006.

8. Rizzolatti, G., Fogassi, L., and Gallese, V. (2001). Neurophysiological mechanisms underlying the understanding and imitation of action. Nat Rev Neurosci 2, 661–670. 10.1038/35090060.

9. Isomura, T., and Nakano, T. (2016). Automatic facial mimicry in response to dynamic emotional stimuli in five-month-old infants. Proc Biol Sci 283. 10.1098/rspb.2016.1948.

10. Soussignan, R., Dollion, N., Schaal, B., Durand, K., Reissland, N., and Baudouin, J.Y. (2018). Mimicking emotions: how 3-12-month-old infants use the facial expressions and eyes of a model. Cogn Emot 32, 827–842. 10.1080/02699931.2017.1359015.

11. Soderkvist, S., Ohlen, K., and Dimberg, U. (2018). How the Experience of Emotion is Modulated by Facial Feedback. J Nonverbal Behav 42, 129–151. 10.1007/s10919-017-0264-1.

12. Oberman, L.M., Winkielman, P., and Ramachandran, V.S. (2007). Face to face: blocking facial mimicry can selectively impair recognition of emotional expressions. Soc Neurosci 2, 167–178. 10.1080/17470910701391943.

13. Conson, M., Ponari, M., Monteforte, E., Ricciato, G., Sara, M., Grossi, D., and Trojano, L. (2013). Explicit recognition of emotional facial expressions is shaped by expertise: evidence from professional actors. Front Psychol 4, 382. 10.3389/fpsyg.2013.00382.

14. Orlandi, A., Zani, A., and Proverbio, A.M. (2017). Dance expertise modulates visual sensitivity to complex biological movements. Neuropsychologia 104, 168–181. 10.1016/j.neuropsychologia.2017.08.019.

15. Christensen, J.F., Gomila, A., Gaigg, S.B., Sivarajah, N., and Calvo-Merino, B. (2016). Dance expertise modulates behavioral and psychophysiological responses to affective body movement. J Exp Psychol Hum Percept Perform 42, 1139–1147. 10.1037/xhp0000176.

16. Calder, A.J., Keane, J., Cole, J., Campbell, R., and Young, A.W. (2000). Facial expression recognition by people with mobius syndrome. Cogn Neuropsychol 17, 73–87. 10.1080/026432900380490.

17. Rives Bogart, K., and Matsumoto, D. (2010). Facial mimicry is not necessary to recognize emotion: Facial expression recognition by people with Moebius syndrome. Soc Neurosci 5, 241–251. 10.1080/17470910903395692.

18. Vannuscorps, G., Andres, M., and Caramazza, A. (2020). Efficient recognition of facial expressions does not require motor simulation. Elife 9. 10.7554/eLife.54687.

19. Giannini, A.J., Tamulonis, D., Giannini, M.C., Loiselle, R.H., and Spirtos, G. (1984). Defective response to social cues in Mobius’ syndrome. J Nerv Ment Dis 172, 174–175. 10.1097/00005053-198403000-00008.

20. Bate, S., Cook, S.J., Mole, J., and Cole, J. (2013). First report of generalized face processing difficulties in mobius sequence. PLoS One 8, e62656. 10.1371/journal.pone.0062656.

21. De Stefani, E., Ardizzi, M., Nicolini, Y., Belluardo, M., Barbot, A., Bertolini, C., Garofalo, G., Bianchi, B., Coude, G., Murray, L., and Ferrari, P.F. (2019). Children with facial paralysis due to Moebius syndrome exhibit reduced autonomic modulation during emotion processing. J Neurodev Disord 11, 12. 10.1186/s11689-019-9272-2.

22. De Stefani, E., Nicolini, Y., Belluardo, M., and Ferrari, P.F. (2019). Congenital facial palsy and emotion processing: The case of Moebius syndrome. Genes Brain Behav 18, e12548. 10.1111/gbb.12548.

23. Kanwisher, N., McDermott, J., and Chun, M.M. (1997). The fusiform face area: a module in human extrastriate cortex specialized for face perception. J Neurosci 17, 4302–4311.

24. Pitcher, D., Japee, S., Rauth, L., and Ungerleider, L.G. (2017). The Superior Temporal Sulcus Is Causally Connected to the Amygdala: A Combined TBS-fMRI Study. J Neurosci 37, 1156–1161. 10.1523/JNEUROSCI.0114-16.2016.

25. Zhang, H., Japee, S., Stacy, A., Flessert, M., and Ungerleider, L.G. (2020). Anterior superior temporal sulcus is specialized for non-rigid facial motion in both monkeys and humans. Neuroimage 218, 116878. 10.1016/j.neuroimage.2020.116878.

26. Ince, R.A., Paton, A.T., Kay, J.W., and Schyns, P.G. (2021). Bayesian inference of population prevalence. Elife 10. 10.7554/eLife.62461.

27. Nicolini, Y., Manini, B., De Stefani, E., Coude, G., Cardone, D., Barbot, A., Bertolini, C., Zannoni, C., Belluardo, M., Zangrandi, A., et al. (2019). Autonomic Responses to Emotional Stimuli in Children Affected by Facial Palsy: The Case of Moebius Syndrome. Neural Plast 2019, 7253768. 10.1155/2019/7253768.

28. Bogart, K., Tickle-Degnen, L., and Ambady, N. (2014). Communicating without the Face: Holistic Perception of Emotions of People with Facial Paralysis. Basic Appl Soc Psych 36, 309–320. 10.1080/01973533.2014.917973.

29. Marneweck, M., Palermo, R., and Hammond, G. (2014). Discrimination and recognition of facial expressions of emotion and their links with voluntary control of facial musculature in Parkinson’s disease. Neuropsychology 28, 917–928. 10.1037/neu0000106.

30. Delicato, L.S. (2020). A robust method for measuring an individual’s sensitivity to facial expressions. Atten Percept Psychophys 82, 2924–2936. 10.3758/s13414-020-02043-w.

31. Yeshurun, Y., Carrasco, M., and Maloney, L.T. (2008). Bias and sensitivity in two-interval forced choice procedures: Tests of the difference model. Vision Res 48, 1837–1851. 10.1016/j.visres.2008.05.008.

32. Darwin, C. (1872). The expression of the emotions in man and animals (J. Murray).

33. Maimon-Mor, R.O., Schone, H.R., Moran, R., Brugger, P., and Makin, T.R. (2020). Motor control drives visual bodily judgements. Cognition 196, 104120. 10.1016/j.cognition.2019.104120.

34. Finzi, E., and Rosenthal, N.E. (2016). Emotional proprioception: Treatment of depression with afferent facial feedback. J Psychiatr Res 80, 93–96. 10.1016/j.jpsychires.2016.06.009.

35. Lundqvist D F.A., öhman A (1998). The Karolinska directed emotional faces (KDEF).. CD ROM from Department of Clinical Neuroscience.

36. Klein, S.A. (2001). Measuring, estimating, and understanding the psychometric function: a commentary. Percept Psychophys 63, 1421–1455. 10.3758/bf03194552.

37. Wichmann, F.A., and Hill, N.J. (2001). The psychometric function: I. Fitting, sampling, and goodness of fit. Percept Psychophys 63, 1293–1313. 10.3758/bf03194544.

38. Keefe, B.D., Villing, M., Racey, C., Strong, S.L., Wincenciak, J., and Barraclough, N.E. (2014). A database of whole-body action videos for the study of action, emotion, and untrustworthiness. Behav Res Methods 46, 1042–1051. 10.3758/s13428-013-0439-6.

39. Cox, R.W. (1996). AFNI: software for analysis and visualization of functional magnetic resonance neuroimages. Comput Biomed Res 29, 162–173. 10.1006/cbmr.1996.0014.

40. Cox, R.W., and Hyde, J.S. (1997). Software tools for analysis and visualization of fMRI data. NMR Biomed 10, 171–178. 10.1002/(sici)1099-1492(199706/08)10:4/5<171::aid-nbm453>3.0.co;2-l.

41. Taylor PA C.G., Glen DR, Rajendra JK, Reynolds RC, Cox RW (2018). FMRI processing with AFNI: Some comments and corrections on “Exploring the Impact of Analysis Software on Task fMRI Results”.

42. VS Fonov, A.E., RC McKinstry, CR Almli, DL Collins (2009). Unbiased nonlinear average age-appropriate brain templates from birth to adulthood. NeuroImage.

43. Kundu, P., Inati, S.J., Evans, J.W., Luh, W.M., and Bandettini, P.A. (2012). Differentiating BOLD and non-BOLD signals in fMRI time series using multi-echo EPI. Neuroimage 60, 1759–1770. 10.1016/j.neuroimage.2011.12.028.

44. Talairach, J., and Tournoux, P. (1988). Co-planar stereotaxic atlas of the human brain : 3-dimensional proportional system : an approach to cerebral imaging (Georg Thieme).

