## Supplemental Material for "Inability to make facial expressions dampens emotion perception"

#### A. Participant Information

| | | **Moebius Syndrome group** | **Control group** |
|---|---|---|---|
| Number of participants | | 24 | 50 |
| Age | | 17 – 64 years | 20 – 63 years |
| Sex | Male<br>Female | 12<br>12 | 27<br>23 |
| Diagnosis | Moebius Syndrome<br>Moebius Syndrome with atypical symptoms[b]<br>Moebius/Poland Syndrome | 20[a]<br>3<br>1 | Healthy volunteer |
| Facial weakness | Bilateral severe facial palsy<br>Bilateral mild facial palsy<br>Unilateral facial palsy | 20<br>2<br>2 | -<br>-<br>- |
| Smile surgery | Bilateral<br>Unilateral | 10<br>1 | -<br>- |
| Horizontal gaze palsy | Bilateral abduction and adduction limitation<br>Bilateral abduction limitation<br>Unilateral abduction limitation | 19<br>3<br>2 | -<br>-<br>- |
| Vertical gaze palsy | Bilateral mild limitation<br>Unilateral limitation | 5<br>2 | -<br>- |
| Eye surgery | Strabismus | 11 | - |
| Intellectual functioning | High average<br>Average<br>Low average<br>Below average | 2[c]<br>14[c], 6[d]<br>1[ce]<br>1[ce] | -<br>50[d]<br>-<br>- |
| Limb anomalies | Transverse terminal limb defects<br>Club feet<br>Brachysyndactyly (short, webbed digits) | 2<br>7<br>2 | -<br>-<br>- |

[a] includes 1 self-reported
[b] atypical symptoms include unilateral facial paralysis and seizures
[c] assessed via Weschler Adult Intelligence Scale
[d] presumptive based on clinical history and interview
[e] two individuals with low to below average IQ performed similar to controls on the static face and dynamic body control tasks, and were thus included in the group analysis

#### B. Moebius Syndrome Collaborative Research Consortium:

**Icahn School of Medicine at Mount Sinai, New York, NY:** Monica Erazo (Department of Genetics and Genomic Sciences, and Department of Neuroscience and Physiology, New York University Grossman School of Medicine), Tamiesha Frempong (Department of Ophthalmology), Ke Hao (Department of Genetics and Genomic Sciences and Icahn Institute for Data Science and Genomic Technology), Ethylin

Wang Jabs (Departments of Genetics and Genomic Sciences, Cell, Developmental and Regenerative Biology, and Pediatrics), Thomas P. Naidich (Departments of Radiology, Neurosurgery, and Pediatrics), Janet C. Rucker (Department of Neurology, and Department of Neuroscience and Physiology, New York University Grossman School of Medicine), Bryn D. Webb (Departments of Genetics and Genomic Sciences and Pediatrics, & Department of Pediatrics, Division of Genetics and Metabolism, University of Wisconsin-Madison, Madison, WI), and Zhongyang Zhang (Department of Genetics and Genomic Sciences and Icahn Institute for Data Science and Genomic Technology).

**Boston Children's Hospital (BCH) and Harvard Medical School (HMS), Boston, MA**: Brenda J. Barry (Department of Neurology. BCH and HMS; Howard Hughes Medical Institute, Chevy Chase, MD), Wai-Man Chan (Department of Neurology, BCH; Howard Hughes Medical Institute. Chevy Chase, MD), Silvio Alessandro DiGioia (Department of Neurology, BCH and HMS), Elizabeth Engle (Departments of Neurology and Ophthalmology, BCH and HMS; Howard Hughes Medical Institute, Chevy Chase, MD); David G. Hunter (Department of Ophthalmology, BCH and HMS), Julie Jurgens (Department of Neurology, BCH and HMS), Arthur Lee (Department of Neurology. BCH and HMS), Sarah E. MacKinnon (Department of Ophthalmology, BCH), Caroline Robson (Department of Radiology, BCH and HMS), Matthew Rose (Departments of Neurology and Pathology, BCH; Department of Pathology, Brigham and Women's Hospital and HMS), and Alan Tenney (Department of Neurology. BCH and HMS).

**National Institutes of Health, Bethesda, MD**: Barbara B. Biesecker (Social and Behavioral Research Branch, National Human Genome Research Institute), Lori L. Bonnycastle (Medical Genomics and Metabolic Genetics Branch, National Human Genome Research Institute), Brian P. Brooks (Ophthalmic Genetics & Visual Function Branch, National Eye Institute), John A. Butman (Radiology and Imaging Sciences, Clinical Research Center), Peter S. Chines (Medical Genomics and Metabolic Genetics Branch, National Human Genome Research Institute), Francis S. Collins (Medical Genomics and Metabolic Genetics Branch, National Human Genome Research Institute and NIH Office of the Director), Flavia Facio (Medical Genomics and Metabolic Genetics Branch, Social and Behavioral Research Branch, National Human Genome Research Institute), Kathleen Farrell (Rehabilitation Medicine Department, Clinical Research Center), Edmond J. FitzGibbon (Ophthalmic Genetics & Visual Function Branch, National Eye Institute), Andrea L. Gropman (George Washington University and Children's National Medical Center, Washington, DC), Elizabeth Hutchinson (Quantitative Medical Imaging Section, National Institute of Biomedical Imaging and Bioengineering, Bethesda, MD), Mina S. Jain (Rehabilitation Medicine Department, Clinical Research Center), Shruti Japee (Laboratory of Brain and Cognition, National Institute of Mental Health), Kelly A. King (Audiology Unit, Otolaryngology Branch, National Institute of Deafness and other Communications Disorders), Tanya J. Lehky (Electromyography Section, National Institute of Neurological Disorders and Stroke), Janice Lee (Office of the Clinical Director, National Institute of Dental and Craniofacial Research), Denise K. Liberton (Office of the Clinical Director, National Institute of Dental and Craniofacial Research), Irini Manoli (Medical Genomics and Metabolic Genetics Branch, National Human Genome Research Institute), Narisu Narisu (Medical Genomics and Metabolic Genetics Branch, National Human Genome Research Institute), Scott M. Paul (Rehabilitation Medicine Department, Clinical Research Center), Carlo Pierpaoli (Quantitative Medical Imaging Section, National Institute of Biomedical Imaging and Bioengineering), Neda Sadeghi (Quantitative Medical Imaging Section, National Institute of Biomedical Imaging and Bioengineering), Joseph Snow (Office of the Clinical Director, National Institute of Mental Health), Beth Solomon (Rehabilitation Medicine Department, Clinical Research Center), Angela Summers (Office of the Clinical Director, National Institute of Mental Health), Amy J. Swift (Medical Genomics and Metabolic Genetics Branch, National Human Genome Research Institute), Camilo Toro (NIH Undiagnosed Diseases Program, Common Fund, National Human Genome Research Institute), Audrey Thurm (Pediatrics and Developmental Neuroscience Branch, National Institute of Mental Health), Carol Van Ryzin (Medical

Genomics and Metabolic Genetics Branch, National Human Genome Research Institute), and Chris K. Zalewski (Audiology Unit, Otolaryngology Branch, National Institute of Deafness and other Communications Disorders).

### C. Signal Detection theory analysis of catch trials in the static face task:

| Task | Emotion | Measure | Controls (M ± SD) | MBS (M ± SD) | **p-value** (between group independent samples t-test) |
|---|---|---|---|---|---|
| Control Task | Fearful | Sensitivity (d') | 3.28 ± 0.36 | 2.95 ± 0.65 | 0.09 |
| | | Criterion (c) | -0.99 ± 0.26 | -1.03 ± 0.28 | 0.68 |
| | Happy | Sensitivity (d') | 2.72 ± 0.44 | 2.80 ± 0.38 | 0.54 |
| | | Criterion (c) | -0.72 ± 0.27 | -0.72 ± 0.19 | 0.98 |
| Emotion Task | Fearful | Sensitivity (d') | 2.78 ± 0.49 | 2.54 ± 0.50 | 0.16 |
| | | Criterion (c) | -0.72 ± 0.27 | -0.39 ± 0.56 | 0.06 |
| | Happy | Sensitivity (d') | 2.82 ± 0.33 | 2.90 ± 0.45 | 0.58 |
| | | Criterion (c) | -0.72 ± 0.16 | -0.60 ± 0.30 | 0.18 |

### D. Statistical analyses for each behavioral task:

* Indicates significance at $\alpha = 0.05$; + indicates Bonferroni corrected for multiple comparisons

#### i. Static Facial Expression Task:

| **Beta Regression Analysis** | | | | | | |
|---|---|---|---|---|---|---|
| **Main effects and Interactions** | **estimate** | **standard error** | **Z value** | **p value** | | |
| Group (Controls, MBS) | -0.16 | 0.04 | 3.92 | 8.72x$10^{-5}$ * | | |
| Task (Control, Emotion) | -0.17 | 0.02 | 6.80 | 1.02x$10^{-11}$ * | | |
| Expression (Fearful, Happy) | 0.21 | 0.02 | 8.70 | 2.0x$10^{-16}$ * | | |
| Task x Group | 0.09 | 0.02 | 3.54 | 0.0004 * | | |
| Task x Expression | -0.05 | 0.02 | 1.84 | 0.066 | | |
| Expression x Group | -0.02 | 0.02 | 0.62 | 0.537 | | |
| Task x Expression x Group | 0.05 | 0.02 | 2.23 | 0.026 * | | |
| **Between Group: Pairwise comparisons** | **Controls** (M ± SD) | **MBS** (M ± SD) | **degrees of freedom** | **t value** | **p value** | **effect size** (Cohen's d) |
| Emotion Task: Happy | 24.2 ± 5.6 | 30.3 ± 6.7 | 3.07 | 3.07 | .0026 *+ | 0.28 |
| Emotion Task: Fearful | 29.2 ± 6.5 | 45.7 ± 13.9 | 7.65 | 7.65 | 2.02x$10^{-14}$ *+ | 0.69 |
| Control Task: Happy | 20.1 ± 5.2 | 23.2 ± 5.6 | 1.64 | 1.64 | 0.10 | 0.15 |
| Control Task: Fearful | 27.8 ± 6.4 | 30.3 ± 6.7 | 1.39 | 1.39 | 0.17 | 0.13 |

#### ii. Dynamic Facial Expression Task:

| **Beta Regression Analysis** | | | | | | |
|---|---|---|---|---|---|---|
| **Main effects and Interaction** | **Estimate** | **Standard error** | **Z value** | **p value** | | |
| Group (Controls, MBS) | -0.27 | 0.06 | 4.66 | 3.11x$10^{-6}$ * | | |
| Task (Control, Emotion) | -0.13 | 0.05 | 2.81 | 0.005 * | | |
| Task x Group | 0.03 | 0.05 | 0.60 | 0.55 | | |
| **Pairwise comparisons** | **Threshold** M ± SD | **Threshold** M ± SD | **Degrees of freedom** | **t value** | **p value** | **Effect size** (Cohen's d) |
| Group: Controls vs. MBS | Controls: 20.7 ± 7.7 | MBS: 31.2 ± 11.2 | 57.5 | 5.89 | 2.12x$10^{-7}$ *+ | 0.78 |
| Task: Control vs. Emotion | Control: 29.2 ± 6.5 | Emotion: 45.7 ± 13.9 | 57.5 | 2.81 | .007 *+ | 0.37 |

| Within Emotion Task: Main effects and Interaction | Estimate | Standard Error | Z value | p value | | |
|---|---|---|---|---|---|---|
| Group (Controls, MBS) | -0.31 | 0.06 | 5.24 | 1.59x10$^{-7}$ * | | |
| Expression (Angry, Fearful) | 0.18 | 0.04 | 4.10 | 4.20x10$^{-5}$ * | | |
| Expression (Angry, Happy) | 0.35 | 0.04 | 8.15 | 3.63x10$^{-16}$ * | | |
| Group x Expression (Angry, Fearful) | -0.12 | 0.04 | 2.89 | 0.004 * | | |
| Group x Expression (Angry, Happy) | 0.08 | 0.04 | 1.84 | 0.07 | | |
| **Between Group: Pairwise comparisons** | **Controls** (M ± SD) | **MBS** (M ± SD) | **Degrees of freedom** | **t value** | **p value** | **Effect size** (Cohen's d) |
| Emotion Task: Happy | 15.3 ± 3.9 | 24.0 ± 6.9 | 79.9 | 4.49 | 2.37x10$^{-5}$ *⁺ | 0.50 |
| Emotion Task: Fearful | 31.3 ± 10.0 | 42.0 ± 12.3 | 79.9 | 4.68 | 1.15x10$^{-5}$ *⁺ | 0.52 |
| Emotion Task: Angry | 24.0 ± 8.0 | 42.8 ± 9.1 | 79.9 | 8.48 | 9.32x10$^{-13}$ *⁺ | 0.95 |
| Control Task | 19.3 ± 8.7 | 28.0 ± 10.8 | 57.5 | 3.65 | 0.0006 *⁺ | 0.48 |

#### iii. Dynamic Body Expression Task:

| Beta Regression Analysis | | | | | | |
|---|---|---|---|---|---|---|
| **Main effects and Interaction** | **Estimate** | **Standard error** | **Z value** | **p value** | | |
| Group (Controls, MBS) | -1.06 | 0.07 | 15.22 | 0.18 | | |
| Task (Control, Emotion) | -0.09 | 0.07 | -1.33 | 2.0x10$^{-16}$ * | | |
| Task x Group | -0.64 | 0.06 | 10.26 | 0.09 | | |
| **Pairwise comparisons** | **Threshold** M ± SD | **Threshold** M ± SD | **Degrees of freedom** | **t value** | **p value** | **Effect size** (Cohen's d) |
| Group: Controls vs. MBS | Controls: 25.0 ± 9.24 | MBS: 29.7 ± 9.34 | 48.4 | 1.49 | 0.14 | 0.21 |
| Task: Control vs. Emotion | Control: 15.1 ± 6.98 | Emotion: 39.6 ± 11.4 | 48.4 | 10.26 | 9.85x10$^{-16}$ *⁺ | 1.48 |
| **Within Emotion Task: Main effects and Interaction** | **Estimate** | **Standard error** | **Z value** | **p value** | | |
| Group (Controls, MBS) | -0.21 | 0.10 | -2.01 | 0.04 * | | |
| Expression (Angry, Fearful) | -0.41 | 0.10 | -4.17 | 3.04x10$^{-5}$ * | | |
| Expression (Angry, Happy) | 0.43 | 0.10 | 4.46 | 8.33x10$^{-6}$ * | | |
| Group x Expression (Angry, Fearful) | -0.12 | 0.10 | -1.19 | 0.23 | | |
| Group x Expression (Angry, Happy) | 0.04 | 0.10 | 0.39 | 0.70 | | |

#### iv. Facial Identity and Expression Matching Task:

| Beta Regression Analysis | Estimate | Standard error | Z value | p value | | |
|---|---|---|---|---|---|---|
| **Main effects and Interactions** | | | | | | |
| Group (Controls, MBS) | 0.75 | 0.16 | 1.02 | 0.31 | | |
| Task (Identity, Expression) | 0.16 | 0.16 | 4.26 | 2.05x10$^{-5}$ * | | |
| Task x Group | 0.30 | 0.07 | 2.14 | 0.03 * | | |
| **Between Group: Pairwise comparisons** | **Controls** (M ± SD) | **MBS** (M ± SD) | **Degrees of freedom** | **t value** | **p value** | **Effect size** (Cohen's d) |
| Identity Task | 60.5 ± 18.6 | 60.2 ± 17.3 | 24.3 | 0.10 | 0.93 | 0.02 |
| Expression Task | 79.1 ± 8.1 | 67.5 ± 8.3 | 24.3 | 3.00 | 0.006 *⁺ | 0.61 |

**E. Bayesian Prevalence analysis of task-based fMRI data:**

We reanalyzed the blocked design fMRI data to extract one beta estimate for each block of the *identity* and *expression* tasks resulting in 16 values per subject for each task condition. Then using a bootstrapped between-task comparison for each subject we determined the proportion of participants in each group that showed the expected effect, i.e., activity during *expression* > *identity* processing. Using these group proportions, we then estimate the Bayesian prevalence difference[26] between the two groups. These results are graphed below. Note the similarity in prevalence patterns between the two groups for rFFA and rpSTS regions but not for rAMG.

<u>For right FFA</u> (maximum a posteriori (MAP) estimate: Controls = 0.23; MBS = 0.30; difference = 0.04):

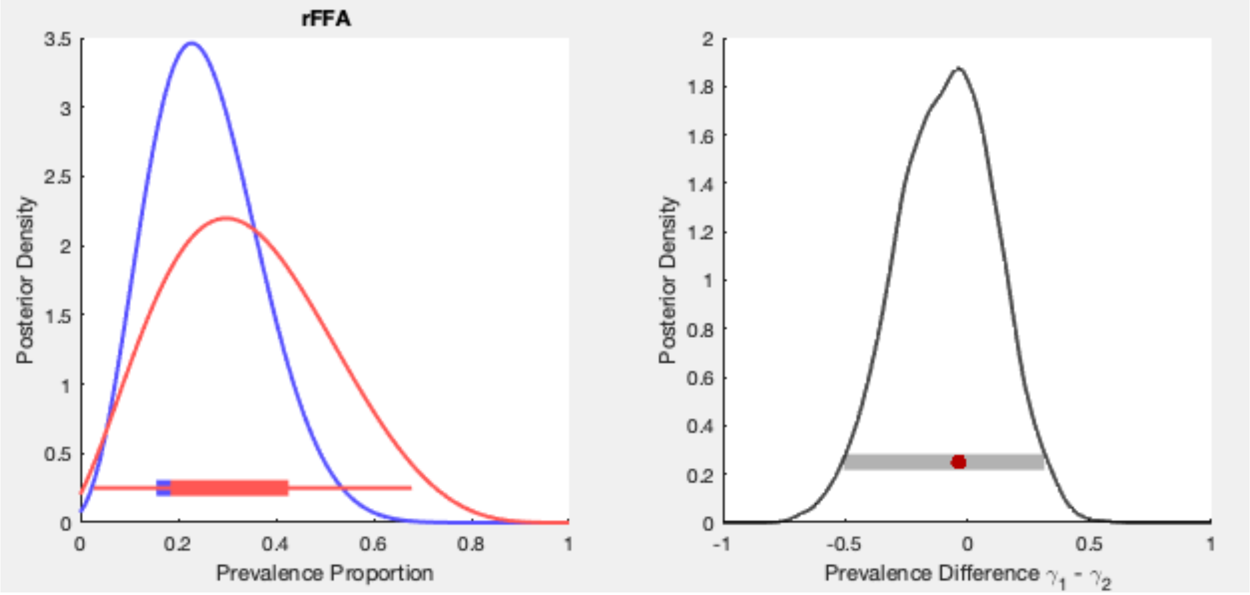

<u>For right pSTS</u> (maximum a posteriori (MAP) estimate: Controls = 0.30; MBS = 0.47; difference = 0.19):

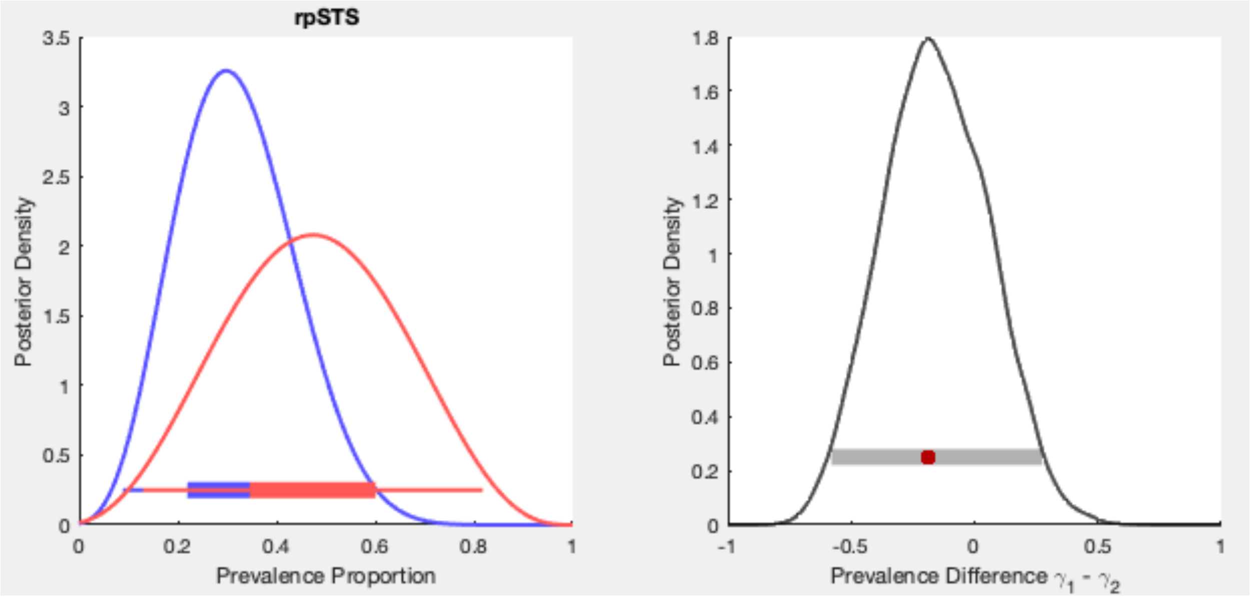

[For right Amygdala] (maximum a posteriori (MAP) estimate: Controls = 0.29; MBS = 0; difference = 0.15):

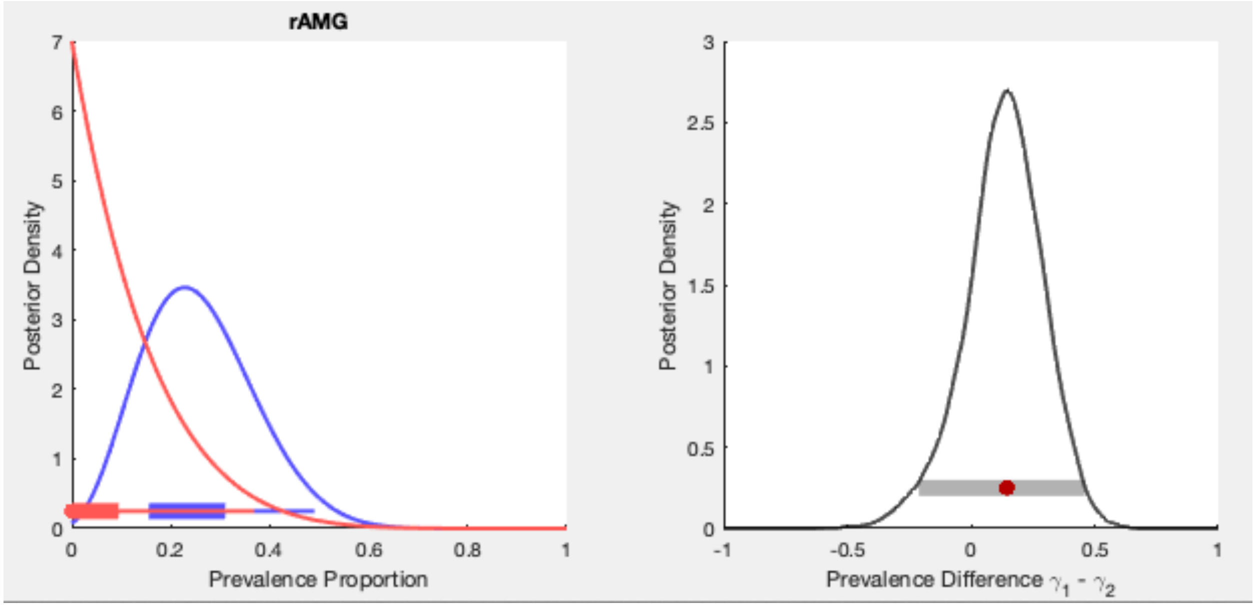
